# A ratiometric ER calcium sensor for quantitative comparisons across cell types and subcellular regions

**DOI:** 10.1101/2024.02.15.580492

**Authors:** Ryan J. Farrell, Kirsten G. Bredvik, Michael B. Hoppa, S. Thomas Hennigan, Timothy A. Brown, Timothy A. Ryan

## Abstract

The endoplasmic reticulum (ER) is an important regulator of Ca^2+^ in cells and dysregulation of ER calcium homeostasis can lead to numerous pathologies. Understanding how various pharmacological and genetic perturbations of ER Ca^2+^ homeostasis impacts cellular physiology would likely be facilitated by more quantitative measurements of ER Ca^2+^ levels that allow easier comparisons across conditions. Here, we developed a ratiometric version of our original ER-GCaMP probe that allows for more quantitative comparisons of the concentration of Ca^2+^ in the ER across cell types and sub-cellular compartments. Using this approach we show that the resting concentration of ER Ca2+ in primary dissociated neurons is substantially lower than that in measured in embryonic fibroblasts.

## Introduction

The endoplasmic reticulum (ER) performs important cellular roles including protein synthesis and glycosylation, lipid homeostasis and calcium regulation (Schwarz and Blower, 2016). As such, the ER emerged as a central hub in stress signaling, relaying the physiological status of ER processes to adaptive transcriptional mechanisms that maintain cellular homeostasis (Hetz et al., 2020). Disruption of ER Ca^2+^ handling can trigger the Unfolded Protein Response/ER stress pathways (Hetz, 2012) and is known to trigger the activation of plasma membrane Ca^2+^ channels (Hogan et al., 2010; Moccia et al., 2015). Additionally, ER Ca^2+^ regulation is critical for a number of cellular processes including contraction of muscle cells (Bers, 2002; Sweeney and Hammers, 2018) and V(D)J recombination in immune cells (Chen et al., 2021). Methods that allow quantitative interrogations of ER Ca^2+^ handling have proven pivotal in discovering novel aspects of ER function. Early work to elucidate ER Ca^2+^ regulation relied on cytosolic calcium changes as a proxy measure of ER Ca^2+^ changes, which were naturally biased for detecting ER Ca^2+^ efflux. Subsequently, probes to directly measure ER Ca^2+^ were developed (Palmer et al., 2004) that were attuned for the concentration of ER Ca^2+^ (∼100 µM), approximately 1000-fold higher than the Ca^2+^ concentration of the cytosol (∼100 nM) (Berridge et al., 2000). We previously created two such ER Ca^2+^ sensors based on the GCaMP6 series with both a large dynamic range and a Ca^2+^ affinity appropriate for the ER. These sensors (ER-GCaMP6-150 and ER-GCaMP6-210) provided measures of ER Ca^2+^ dynamics and levels in ER compartments as small as those found in the axons of hippocampal neurons (de Juan-Sanz et al., 2017). This provided some of the first measurements of ER lumen Ca^2+^ influx, which were temporally locked with neuronal activity. Although suitable for looking at dynamics, quantitative comparisons of signals derived from these reporters across different cells or subcellular locations was laborious as it relied on permeabilizing the cells and comparing the resting fluorescence with the maximal saturated fluorescence signal. This approach corrects for the expression level of the reporter since signals are compared internally to the saturated value. Here we report a modification of our original ER-GCaMPs where we fused the HaloTag protein (Los et al., 2008) to ER-GCaMP6 to create a ratiometric ER-GCaMP6 sensor that alleviates the need to correct for sensor expression levels. Instead, expression levels can be inferred directly from the HaloTag channel signal allowing for rapid and faithful determination of resting ER Ca^2+^ levels in cells. Although red fluorescent proteins spectrally separated from GFP would in principle be suitable for this purpose, the differential susceptibility of coral-based red fluorescent proteins to degradation compared to GFP (Katayama et al., 2011) led to an unbalanced accumulation of the red compared to the green proteins in chimeric constructs with ER-GCaMP. In contrast, chimeric sensors fused with Halo showed no aberrant accumulation. Using this sensor, we show that neurons have significantly lower ER Ca^2+^ levels than mouse embryonic fibroblasts but reveal a large variation in ER Ca^2+^ levels across cell types as well as across cells of the same type. This new sensor (ER-Halo-GCaMP6-150) provides a powerful tool to study ER Ca^2+^ regulation across many cell types and may also facilitate the discovery of new ER Ca^2+^ regulators.

## Results

### Development of a Ratiometric ER-GCaMP

We previously created an ER Ca^2+^ indicator based on the GCaMP6 series (ER-GCaMP6-150) that retained the large dynamic range of GCaMP6 fluorescence change upon Ca^2+^ binding, but whose affinity was weakened by a factor of ∼1000X (de Juan-Sanz et al., 2017). When targeted to the lumen of the ER this sensor was suitable for examining Ca^2+^ levels within the ER. In order to interpret the absolute intensity of the sensor, a necessity for making comparisons between different conditions or cell types, one must deconvolve the signal contribution from sensor expression level from signal arising from variations in [Ca^2+^]. In principle many fluorescent probes have a true isosbestic point, i.e. a wavelength of excitation or emission that is insensitive to changes in ligand occupancy, that could be used to account for expression. In practice for GFP-based sensors such as GCaMPs, the isosbestic point is often in the very blue shifted part of the excitation spectrum (Dana et al., 2019), a region where autofluorescence excitation can dominate the signal. An alternate approach is to fuse the GFP-based sensor to a suitably spectrally separated red fluorescent protein, creating a 2-color chimeric construct, where the ratio of the fluorescence arising from the two components effectively normalizes the signal for expression level. We initially screened candidate chimeric constructs by expressing them in neurons and imaged using sequential illumination in the two channels. We discovered that when ER-GCaMP6 was fused to mRuby3 (Figure 1A) and expressed in rat hippocampal neurons, numerous puncta were apparent in the mRuby channel that were not as strong in the ER-GCaMP channel (Figure 1B, orange arrows). This was likely because coral-based proteins such as mRuby are resistant to acid-protease based degradation (Katayama et al., 2011). The mRuby therefore survives much longer in the degradative pathway, creating an artificial imbalance in the green and red signal. Such an indicator would not allow for accurate reporting of ER Ca^2+^ levels, as the mRuby fluorescence was not an accurate measure of functional GCaMP expression. We therefore undertook a different strategy in which we fused the ER-GCaMP6-150 to the HaloTag protein to create the ER-Halo-GCaMP6-150 (Figure 1C). HaloTag is a bacterial dehalogenase that has been modified to covalently bind molecules containing a haloalkane ligand (Los et al., 2008). We leveraged the creation of the bright JaneliaFluor (JF) dyes with a HaloTag ligand for the normalization channel of our ratiometric probe (Grimm et al., 2017a, 2017b). Expression of the ER-Halo-GCaMP-150 in rat hippocampal neurons and subsequently loaded with the far-red JaneliaFluor dye JF_635_ revealed that the two fluorescent channels were much more balanced, without the appearance of puncta as seen with mRuby (Figure 1D).

**Figure 1.**
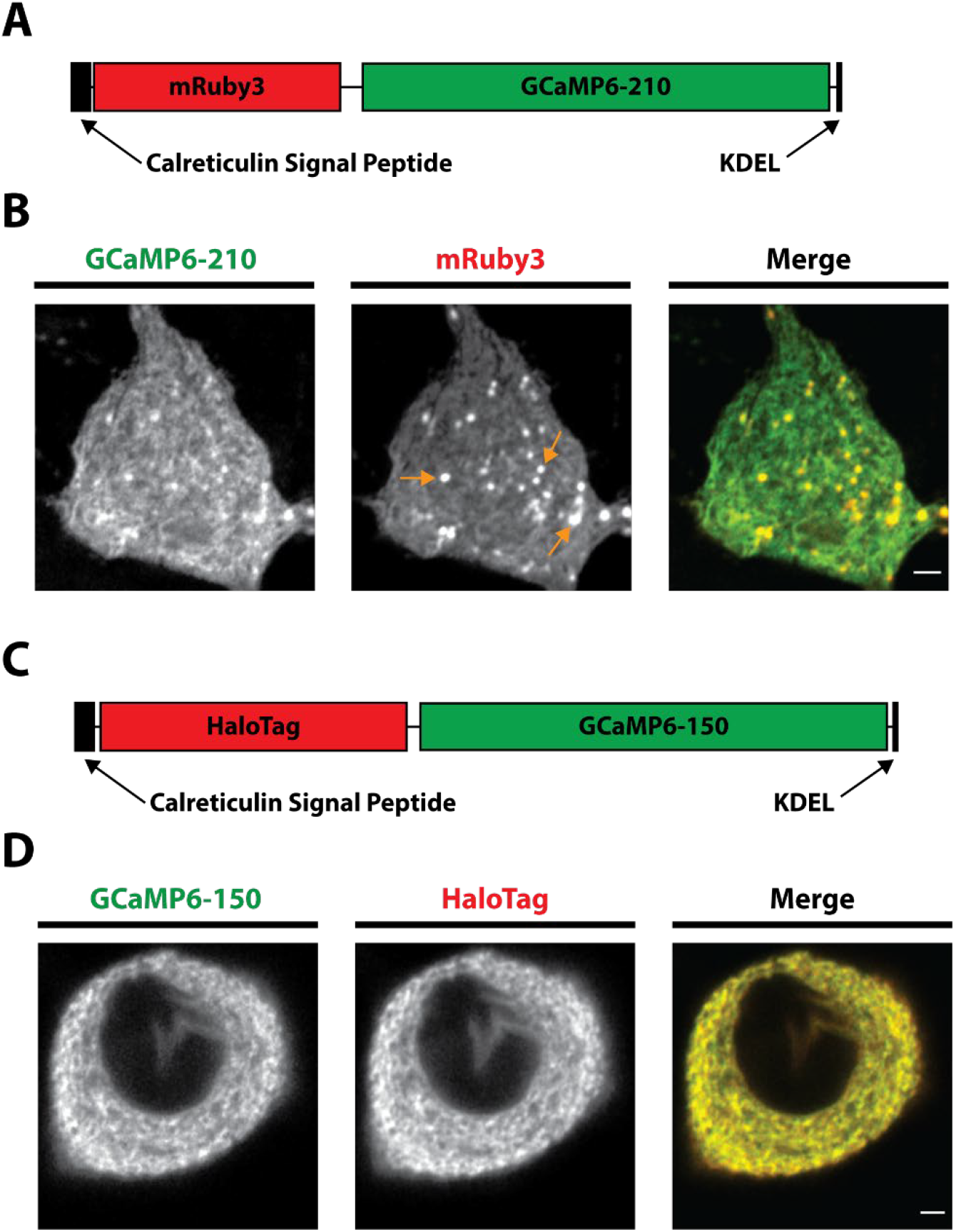
Development of the ER-Halo-GCaMP6-150. **A)** Schematic of the ER-mRuby3-GCaMP6-210. Calreticulin Signal Peptide and KDEL retention sequences are for ER targeting, GCaMP6-210 is the low affinity GCaMP. **B)** Airy scan images showing the expression of the ER-mRuby3-GCaMP6-150 in a hippocampal cell body (GCaMP6-210 - green, mRuby3 – red). Orange arrows mark examples of puncta that are thought to be the sensor undergoing degradation. Scale bar is 2 µm. **C)** Schematic of the ER-Halo-GCaMP6-150. Calreticulin Signal Peptide and KDEL retention sequences are for ER targeting, GCaMP6-150 is the low affinity GCaMP and HaloTag is the bacterial dehalogenase. **D)** Airy scan images showing the expression of the ER-Halo-GCaMP6-150 in a hippocampal cell body (GCaMP6-150 - green, HaloTag – red). HaloTag is loaded with JF-635. Scale bar is 2 µm.

### Ratiometric ER-GCaMP6 calibration

The addition of the HaloTag protein to these sensors could in principle alter the sensor’s binding of Ca^2+^ ions. To characterize the Ca^2+^ binding properties of the ER-Halo-GCaMP6-150, recombinant ER-Halo-GCaMP6-150 was expressed in bacteria, purified and the fluorescence was monitored as a function of Ca^2+^ ion concentration ([Ca^2+^]). The resulting quantitative relationship between [Ca^2+^] and the resulting fluorescence was fit to the generalized Hill Equation, as previously described (de Juan-Sanz et al., 2017). These measurements showed that the new chimeric sensor had only modest changes in the binding parameters compared to the original sensor (Figure 2 A, B, E).

**Figure 2.**
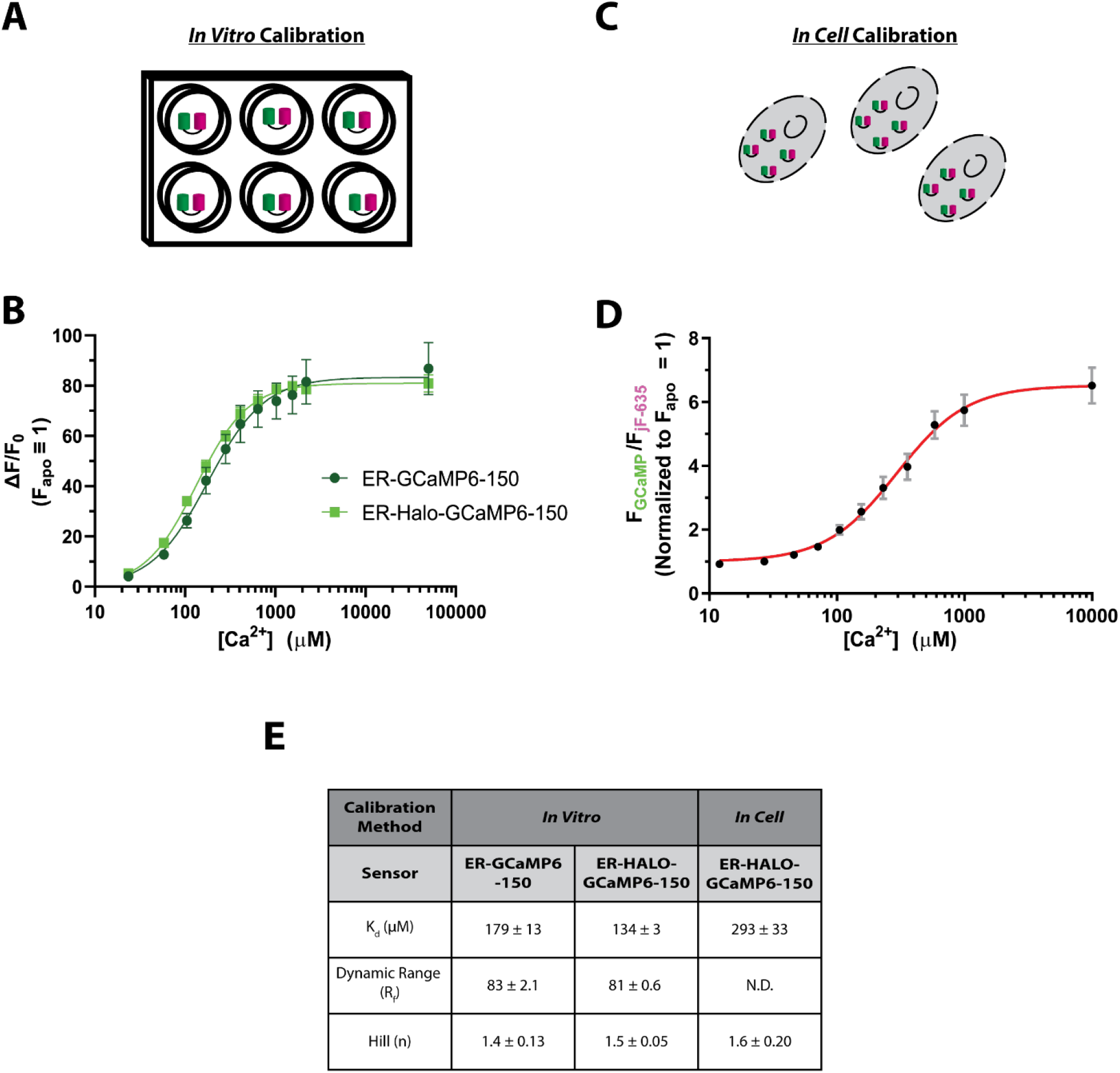
Calibration of the ratiometric ER-GCaMP. **A)** Schematic of the *in vitro* calibration process **B)** Binding curve of the *in vitro* calibration fit to the Hill Equation (ER-GCaMP6-150: n = 4, ER-Halo-GCaMP6-150: n = 4, error bars are ± SEM). **C)** Schematic of the *in cell* calibration process **D)** Binding curve of the *in cell* calibration in rat cortical somas fit to the Hill Equation (ER-Halo-GCaMP6-150: n = 5, error bars are ± SEM). **E)** Table comparing binding characteristics of the Hill Equation from the fits in B) and D). N.D. not determined.

The environment of an indicator expressed in cells is much more complex than that of the purified protein and may impact Ca^2+^ binding properties. To enable accurate measurements of ER Ca^2+^ within cells we determined the binding properties of the ER-GCaMPs using Ca^2+^ titration in permeabilized neurons. Primary dissociated rat cortical neurons previously transfected with ER-Halo-GCaMP6-150 were permeabilized with digitonin (see methods) and the fluorescence monitored during the perfusion of multiple buffered solutions with different [Ca^2+^]. For each [Ca^2+^], ER fluorescence at cell somas for both the GCaMP6-150 and the Halo protein bound to JF_635_ was measured (Figure 2 C, D). The data were then fit to a generalized Hill Equation to determine the relevant binding parameters. As has previously been reported, the *in-cell* binding parameters were somewhat different than those determined *in-vitro*, in particular the binding affinity (K_d_) was weakened by ∼ 120%, but still within a range suitable for measurements of ER Ca^2+^.

### Measurement of ER Ca^2+^ in neurons

Most approaches used to estimate the concentration of Ca^2+^ in cells use probes that themselves are Ca^2+^ buffers. This buffering can potentially influence both how the cell handles Ca^2+^ dynamics, and in a sufficiently isolated compartment, even the resting value free Ca^2+^ (McMahon and Jackson, 2018). One advantage of using a ratiometric indicator is the ability to look for correlations in the reported Ca^2+^ level, i.e. the ratiometric signal, with the expression level of the reporter given by the intensity of the inert channel, in this case Halo-JF_635_. When the ER-Halo-GCaMP6-150 was expressed in rat hippocampal neurons (Fig. 3 A), these measurements showed that when one includes a very wide range of expression levels (∼120 fold), there is a clear inverse correlation between ER Ca^2+^ and expression of the reporter (R^2^ = 0.22) (Fig. 3 B). However, if one restricts observations to the lower 20% of the range of expression (which is where 63% of cells measured landed) the correlation is very weak (R^2^ = 0.08), This suggests that except for in cases of very high ER-Halo-GCaMP6-150 expression levels the buffering capability of the indicator is not a significant concern for accurate quantitation of ER Ca^2+^.

**Figure 3.**
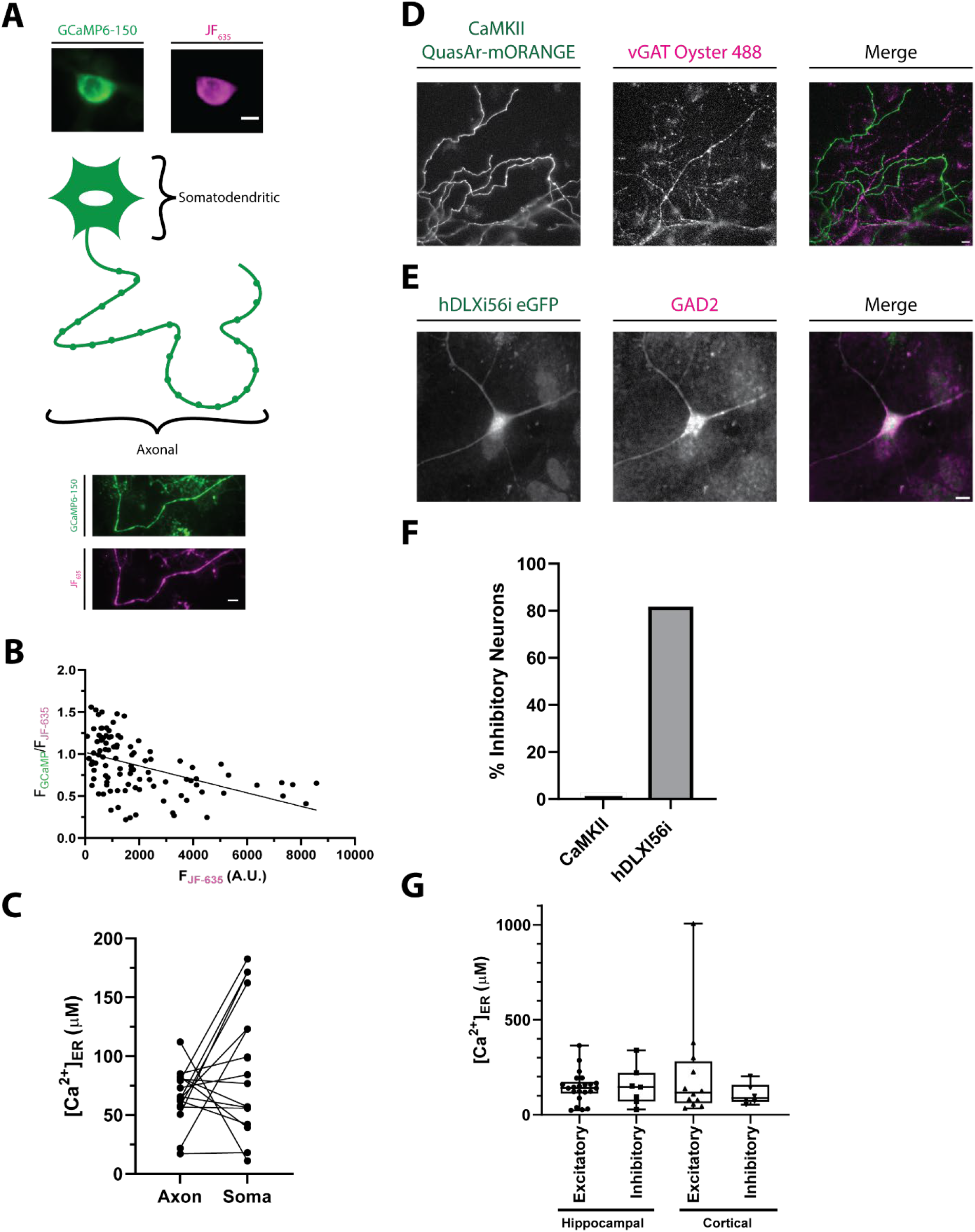
Comparisons of neuronal ER Ca^2+^. **A)** Schematic with sample widefield images of the ER-Halo-GCaMP6-150 loaded with JF_635_. The somatic (top) and axonal (bottom) ER compartments that were compared in B) are highlighted. (ER-GCaMP6-150 is green and the JF_635_ is magenta). Scale bar is 10 µm. **B)** Plot of the ratio of the ER-GCaMP6-150 signal to the JF-Halo bound fluorescence signal as a function of the total JF_635_ Halo-bound fluorescence signal measured over a large population of individual hippocampal cell somas. (n = 101 cells). The expression level of the indicator is weakly correlated with the ratio reporting the ER Ca^2+^ level as reported by a linear fit (line, R^2^ = 0.22). **C)** Comparison of ER [Ca^2+^] in the soma and axonal regions of the same neurons. (n = 16, n.s. p > 0.05, Welch’s t-test) **D)** Representative image showing the specificity of the CaMKII promoter for excitatory neurons. Neurons are transfected with QuasAr-mORANGE under the CaMKII promoter. Following stimulation in the presence of a luminal vGAT antibody, inhibitory nerve terminals are labeled. Scale bar is 10 µm. **E)** Representative immunostaining showing the specificity of the hDLXI56i enhancer for inhibitory neurons. Neurons are transfected with eGFP under the hDLXI56i enhancer which can be recognized by a GFP antibody. GAD2 labels inhibitory neurons. Scale bar is 10 µm. **F)** Quantification of the percentage of inhibitory neurons for the CaMKII promoter (1/36 cells, 2.8%) and the hDLXI56ienhancer (18/22 cells, 81.8%). **G)** Comparison of ER Ca^2+^ levels between inhibitory and excitatory neurons of both the cortex and hippocampal regions of the rat brain. (Excitatory hippocampal n = 22; inhibitory hippocampal n = 7; excitatory cortical n = 12; inhibitory cortical: n = 6; all comparison n.s. p > 0.05, Kruskal-Wallis)

The ER is a continuous structure throughout neurons, but while the somatodendritic compartments contain both smooth and rough ER, the axon contains only smooth ER (Wu et al., 2017) The unique roles of somatodendritic and axonal regions of a neuron may lead to differences in ER Ca^2+^ regulation in these compartments. For example, in response to electrical activity, axonal ER shows a net uptake of Ca^2+^ (de Juan-Sanz et al., 2017) whereas dendrites show a net decrease in ER Ca^2+^ (Hirabayashi et al., 2017). To determine if axonal and somatodendritic ER have different resting ER Ca^2+^ levels, we expressed ER-GCaMP6-Halo-150 in dissociated rat hippocampal neurons and measured ER Ca^2+^ levels of the soma and axon from the same neuron (Fig. 3A). While Ca^2+^ levels were variable between compartments and cells, we observed no consistent difference in ER Ca^2+^ levels between the axon and soma (Fig. 3C). It is interesting to note that the variability of resting somatic ER Ca^2+^ across a population (55 µM) was significantly larger than that in axons (23 µM) (F-test, p = 0.002), suggesting that the axonal region is under stricter feedback regulation.

Beyond compartmentalization, neurons also have many different subtypes. These various subtypes differ in their morphology, function, and activity levels (Zeng and Sanes, 2017) which could also impact resting ER Ca^2+^ levels. To investigate this question, we measured ER Ca^2+^ levels in excitatory and inhibitory neurons in both the hippocampus and the cortex. To target excitatory neurons, we expressed the ER-Halo-GCaMP-150 under the CaMKII promoter (Dittgen et al., 2004). To validate the specificity of this promoter in primary hippocampal culture we expressed QuasAr-mORANGE under this promoter. Neurons were stimulated with 1000 action potentials (AP), 10 Hz in the presence of a vGAT luminal antibody tagged with Alexa Fluor 488, which will label all inhibitory terminals (Fig 3D). Of the 36 cells examined using this protocol, 35 were unambiguously vGAT-negative indicating the CaMKII promoter enriches for excitatory neurons (Fig. 3F). To target inhibitory neurons, we used the hDLXI56i enhancer construct with a minimal beta-globin promoter to drive expression in interneurons (Dimidschstein et al., 2016; Mich et al., 2021; Zerucha et al., 2000). We verified the specificity of the hDLXI56i enhancer in our primary neuron cultures by expressing GFP under this enhancer followed by retrospective immunostaining of GFP and Glutamate Decarboxylase 2 (GAD2) an enzyme whose expression is specific to inhibitory neurons (Fig. 3E). We found that 82% of cells expressing the hDLXI56ieGFP were also positive for GAD2 expression, indicating that the hDLXI56i enhancer strongly restricts expression to inhibitory neurons in our primary neuronal culture system (Fig. 3F). Comparison of ER Ca^2+^ values from these two distinct cell populations derived from a combination of primary hippocampal or cortical cultures did not reveal any consistent differences (Fig. 3G). Notably however, we did observe a large variation in Ca^2+^ levels between cells but the extent of this variation was similar in the two cell groups.

### Comparison ER Ca^2+^ levels in neuronal and a non-neuronal cell type

As excitable cells, neurons have unique constraints on calcium regulation that could cause differences in ER Ca^2+^ regulation compared to non-excitable cells. Indeed, the major regulators of ER Ca^2+^ influx and efflux have cell type specific isoforms (des Georges et al., 2016; Fan et al., 2015; Periasamy and Kalyanasundaram, 2007; Zalk et al., 2015). We therefore chose to profile ER Ca^2+^ levels in a non-excitable cell type, Mouse Embryonic Fibroblasts (MEFs), and compared their ER Ca^2+^ levels to that of primary rat hippocampal neurons (Figure 4A). As with primary neurons, we also observe an inverse correlation between ER Ca^2+^ and expression level (R^2^ = 0.17) in MEFs but this effect is minimized when examining only the lower 20% of the expression range (which contained 90% of the data, R^2^ = 0.09) (Fig 4B). As with primary neurons, we observed a large range of ER Ca^2+^ levels in the MEF population however, but comparing the cumulative distribution of ER Ca^2+^ values measured over a large population of cells shows that MEFs have a significantly elevated ER Ca^2+^ compared to neurons (Fig 1D, ^***^ p < 0.001, Kolmogorov-Smirnov).

**Figure 4.**
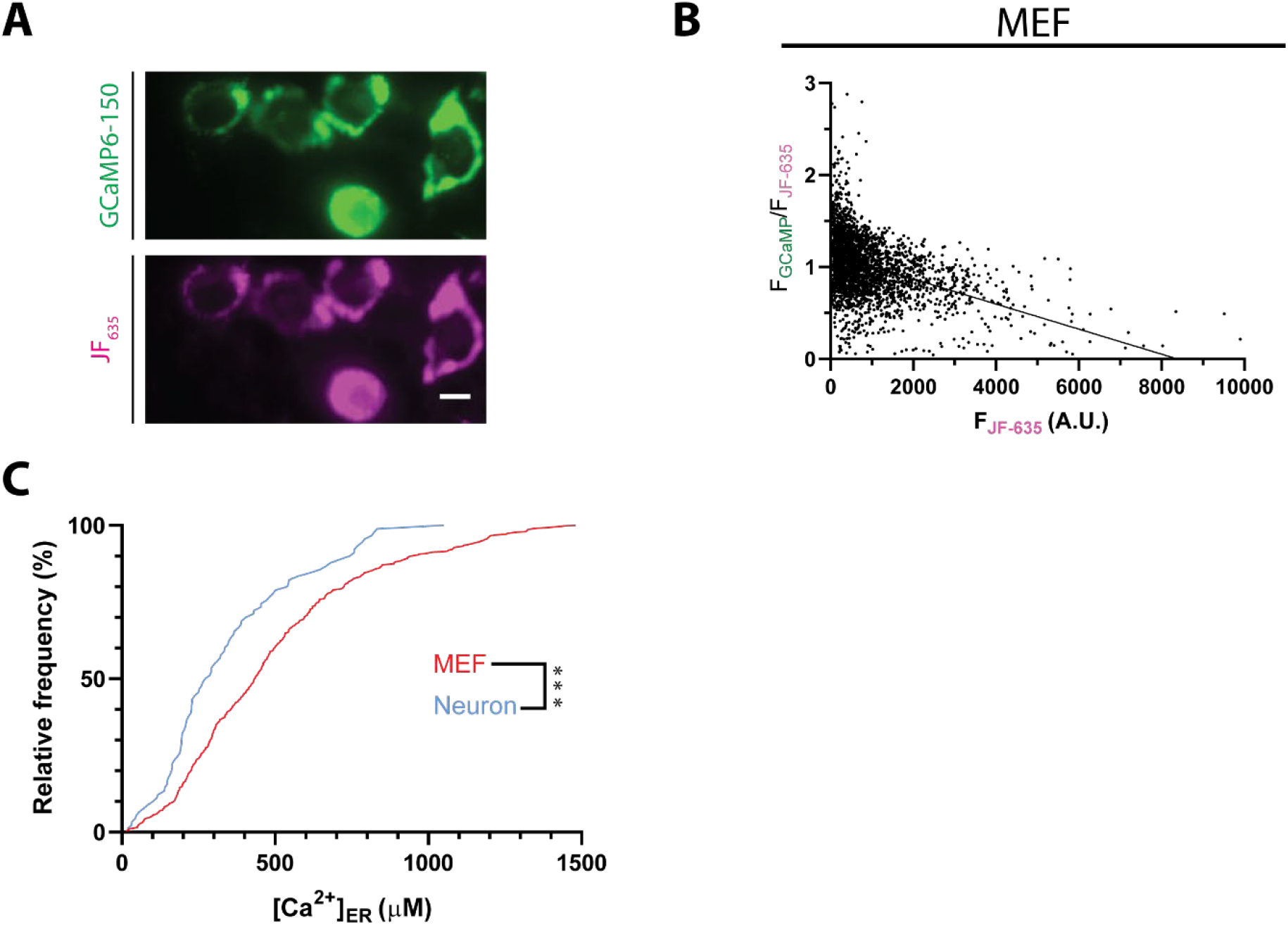
Comparison of neuronal and MEF ER Ca^2+^ levels. **A)** Sample widefield images of a Mouse Embryonic Fibroblasts (MEFs) expressing the ER-Halo-GCaMP6-150 (ER-GCaMP6-150 is green and the JF_635_ is magenta). Scale bar is 10 µm. **B)** Plot of the ratio of the ER-GCaMP signal to the JF-Halo bound fluorescence signal as a function of the total JF Halo-bound fluorescence signal measured over a large population of individual MEF cells. (n = 3,154 cells). The expression level of the indicator is weakly correlated with the ratio reporting the ER [Ca^2+^] level as reported by a linear fit (line, R^2^= 0.17). **C)** Cumulative frequency plots of rat hippocampal neuron soma and mouse embryonic fibroblast (MEF) ER Ca^2+^ values. (Neuron: n = 90, Mean: 345 ± 24 µM; MEF: n = 447, Mean = 492 ± 15 µM ^***^ p < 0.001, Kolmogorov-Smirnov).

## Discussion

Here we developed an approach to ease the ability to make quantitative measurements of ER Ca^2+^ across cells by making use of a chimeric reporter composed of an ER-targeted GCaMP6-based reporter module fused to a HaloTag, which when bound to a fluorescent Halo ligand provides a means to separately normalize the total GCaMP6 signal for expression of the reporter. In general, there are three ways to extract signal from a reporter that is independent of reporter expression. One, if available, is to make measurements at an isosbestic wavelength that by definition should be independent of the relevant analyte. A second is to use a reporter that is optimized for changes in the fluorescence lifetime upon binding the analyte (Koveal et al., 2020), and is therefore independent of expression level itself. As the measured fluorescence lifetime arises from a mixture of analyte bound states, the extracted fluorescence time course reflects the sum of at least two exponential processes, and the parameters are deconvolved from a multi-exponential fit. The third, implemented here, uses a chimeric reporter where one channel, since it is derived from an analyte-insensitive fluorescent protein, serves as a pure measure of expression of the protein.

Creation of such a fusion protein did affect the binding properties of our indicator but still with an appropriate range for ER Ca^2+^ measurements. However, we also observed a difference in binding properties between the purified protein and the indicator when expressed in cells. ER Ca^2+^ indicators used here are a fusion protein containing GFP, calmodulin and the myosin light chain (de Juan-Sanz et al., 2017). Ca^2+^ binds to the EF-hands of calmodulin, leading calmodulin to bind myosin light chain which causes an increase in GFP fluorescence. While the myosin light chain present within the ER-Halo-GCaMP6-150 is in close proximity to the sensor’s calmodulin, making it a likely binding partner, calmodulin is known to interact with many proteins within a cell (Hoeflich and Ikura, 2002). When expressed in cells, calmodulin has access to many of these potential binding partners causing a lower sensor Ca^2+^ affinity when expressed in cells compared to purified protein (Whitaker, 2010). This expectation is in line with the *in cell* calibration having a higher K_d_ (293 µM) than the one found from purified protein (179 µM) (Figure 2). For accurate reporting of Ca^2+^ levels it will be important to measure the binding properties of the indicators in the cell type of interest with the conditions used for experiments.

One advantage of having a readily available readout of the amount of reporter in a given experiment is that it provides straightforward approach to examine whether expression levels impact the reported ER Ca^2+^ values. As our expression levels cover a wide dynamic range (∼100 fold range of expression) we were able to determine a weak negative correlation of expression with the reported resting ER Ca^2+^ and expression levels in both primary neurons and MEFs (Fig 3B. Fig.4B). For practical purposes this inverse correlation is quite minimal however if one restricts the observations to the cells in the lower expression ranges.

Even though the ER is a continuous structure in neurons (Wu et al., 2017), variations were observed between compartments in the ER. It is known that ER Ca^2+^ levels change in response to activity (de Juan-Sanz et al., 2017; Hirabayashi et al., 2017), but how ER Ca^2+^ changes from long-term persistent activity is unknown. Given that ER Ca^2+^ levels dynamically track electrical activity (albeit with a many second time constant), the variability in ER Ca^2+^ levels could be a reflection of the chronic history of activity of the neuron.

Our probe revealed a significant difference in resting ER Ca^2+^ in the non-excitable cell type we examined (MEFs) compared to primary neurons and suggests that this tool will prove valuable in dissecting the molecular pathways of ER Ca^2+^ handling in different cell types. It will be interesting to deploy our ratiometric indicator in muscle and immune cells where ER Ca^2+^ is critical for their function (Lilliu et al., 2021; Trebak and Kinet, 2019). The principles outlined here will provide a basis for use of this indicator in these other cell types and will provide a rapid method to quantify ER Ca^2+^. We expect that this indicator will also be useful in screens as our ratiometric indicator could be used to quantify ER Ca^2+^ levels in sorted cells, providing a high throughput method to identify novel regulators of ER Ca^2+^ regulation such as the unknown ER Ca^2+^ leak pathway (Burgess et al., 1984; Hofer et al., 1996; Thastrup et al., 1990). We believe that our newly developed ratiometric indicator will be valuable in future studies where accurate quantification of ER Ca^2+^ concentrations are critical to understanding biological processes.

### Reagent and Data Availability

Reagent requests will be fulfilled by the lead contact (T.A.R.). The CaMKII ER-Halo-GCaMP6-150 and hDLXI56i ER-Halo-GCAMP6-150 were deposited at Addgene with ID numbers 216316 and 216317 respectively. he data are available from the corresponding author upon reasonable request

## Materials and Methods

### Animals

All neuronal experiments utilized wild-type rats of the Sprague-Dawley strain (Charles River code 400, RRID: RGD_734476). These experiments were conducted in accordance with approved protocols by the Weill Cornell Medicine IACUC. P0-P2 rat pups of mixed gender were sacrificed for further dissection of either the hippocampus or cortex.

### Primary neuronal culture

Hippocampal or cortical neurons were plated on poly-ornithine coated coverslips, transfected via calcium phosphate 6-8 day after plating, and imaged 14-21 days after plating as previously described (Farrell et al., 2023; Ryan, 1999). Neurons were maintained in media composed of MEM (Thermo Fisher 51200038), 0.6% glucose, 0.1 g/L bovine transferrin (Millipore Sigma 616420), 0.25 g/L insulin (Millipore Sigma I6634), 0.3 g/L GlutaMAX (Thermo Fisher 35050061), 5% fetal bovine serum (R&D Systems S11550), 2% homemade NS-21 (Chen et al., 2008) and 4 µM cytosine β-D-arabinofuranoside (Millipore Sigma C6645). Neurons were incubated at 37 °C in a 95% air/5% CO_2_ incubator until imaging.

### MEF Cultures

Mouse Embryonic Fibroblasts (a gift of Yueming Li, MSKCC) were cultured in media composed of DMEM (Thermo Fisher 11965092), Penicillin-Streptomycin/L-glutamine (Thermo Fisher 10378016) and 5% Fetal Bovine Serum (R&D Systems S11550). Cells were transfected using Lipofectamine 3000 (Thermo Fisher L3000015) following the manufacturers protocol with 1.25 µg of DNA/plasmid per 35 mm culture plate. Cells were incubated at 37 °C in a 95% air/5% CO_2_ incubator until imaging.

### Plasmid Constructs

To construct the ER-GCaMP fused to mRuby3, mRuby3 was PCR amplified from the pKanCMV-mRuby3-18aa-actin (Bajar et al., 2016) (Addgene #74255). CMV ER-GCaMP6-210 was restriction digested with AgeI-HF and combined with the PCR amplified mRuby3 via ligation. To construct the ER-GCaMPs fused to Halo, an ER targeted Halo (ER-Halo) was synthesized (Gene Art, Thermo Fisher) using the same ER targeting strategy for the ER-GCaMPs (de Juan-Sanz et al., 2017). The ER-Halo and the ER-GCaMP6-150 (Addgene #86918) under a CamKII promoter were both restriction digested with AgeI and the HALO protein was ligated with Quick Ligase (NEB) into the ER-GCaMP yielding the CamKII ER-Halo-GCaMP6-150. To construct the hDLXI56i eGFP, CMV eGFP-synapsin1a (Chi et al., 2001) was restriction digested with BamHI and BglII. Sticky ends were ligated together with Quick Ligase to remove synapsin1a, making CMV eGFP. The hDLXI56i enhancer was PCR amplified from a hDLXI56i mSynaptophysin-pHluorin plasmid. The CMV eGFP plasmid was restriction digested with AgeI-HF and AseI. The hDLXI56i enhancer was combined with digested CMV eGFP using NEBuilder HiFi DNA Assembly Master Mix (HiFi). To construct the hDLXI56i ER-HALO-GCaMP6-150, the hDLXI56i mSynaptophysin-pHluorin construct was cut with XbaI and AflII. The ER-Halo-GCaMP6-150 was PCR amplified and the resulting fragment was combined with the hDLXI56i backbone with HiFi (NEB). The hDLXI56i enhancer was originally obtained from CN1851-rAAV-hI56i-minBglobin-iCre-4×2C-WPRE3-BGHpA (Graybuck et al., 2021) (Addgene #164450).

### Labeling with jF Dyes

To label cells expressing the ER-Halo-GCaMP6-150, 500 nM JF_635_ containing the HaloTag ligand (Grimm et al., 2017a, 2017b)was added to culture media for 30 minutes. Cells were then washed three times for 10 minutes in fresh culture media.

### Confocal imaging

Neurons were imaged live on a Zeiss LSM 880 with AiryScan in the Weill Cornell Microscopy and Image Analysis Core Facility. Samples were imaged with a 40X, 1.3 NA oil objective. AiryScan deconvolution was performed in the Zeiss Zen software to improve resolution and reveal the ER morphology.

### Live Imaging of Neurons

Imaging experiments were performed on a custom-built laser illuminated epifluorescence microscope with two Andor iXon+ cameras (model DU-897 E-CS0-#BV) and TTL controlled Coherent OBIS 488 nm, 561 nm, and 637 nm lasers. Images were acquired through a 40X, 1.3 NA Fluar Zeiss Objective. Coverslips were mounted in a laminar flow perfusion chamber and perfused at 0.1 mL/min with Tyrodes buffer containing (in mM) 119 NaCl, 2.5 KCl, 1.2 CaCl_2_, 2.8 MgCl_2_, 25 HEPES, 30 glucose, supplemented with 10 µM 6-cyano-7-nitroquinoxabile-2, 3-dione (CNQX) and 50 µM D,L-2-amino-5-phosphonovaleric acid (APV) at pH 7.4. Temperature was controlled at 37 °C using a custom-built heating element. Action potentials were evoked with 1 ms pulses creating field potentials of ∼10 V/cm via a platinum-iridium electrode. Timing of stimulation was controlled by custom-built Arduino and python programs.

For CaMKII promoter validation experiments, neurons expressing CaMKII QuasAr-mORANGE were stimulated 1000 AP, 10 HZ in the presence of 1:100 anti VGAT luminal domain polyclonal rabbit antibody conjugated to Alexa Fluor 488 (Synaptic Systems 131 103C2). Following washout, terminals showing antibody signal were deemed inhibitory.

### Live Imaging of MEF Cells

Imaging was performed on the same set up described for neurons. Tyrodes buffer did not contain APV and CNQX.

### Protein Expression and Purification

Protein expression was initiated by transforming ∼80 ng of each DNA plasmid into 50 µL of XL-1 Chemically Supercompetent Cells (Agilent Technologies, 200236) at 42 °C for 30 seconds followed by 5 minutes of incubation on ice. 10 µL of the bacteria were plated onto LB agar with 60 µg/mL of ampicillin and incubated overnight at 37 °C. A single colony was expanded in 500 mL of complete ZY media (∼450 mL ZY media, 10 mL of 50X salts, 10 mL of 50X 5052, 1 mL of 1M MgSO_4_, 0.1 mL of 1000x trace minerals, and 0.5 mL of 1000X ampicillin [100 µg/mL]) overnight at 30 °C shaking. The culture was centrifuged at 2500 x g for 10 min, and the pellet resuspended in 30 mL of 1M NaCl in PBS, and frozen at -20°C. After thawing in warm water, the sample was sonicated for 5 minutes on ice at Power 10 (Fisher Brand, CL-18). The lysate was spun down at 8000 x g for 10 min to pellet the cellular debris followed by an additional spin for 45 minutes at 35000 x g.

For protein purification, the supernatant was run over a nickel-based HisTrap HP column (Cytiva, 17524801) on a Bio-Rad BioLogic LP system. The column was generated, and the lysate loaded as per manufacturer’s instructions. The protein was eluted off in fractions using 0.2 M imidazole PBS. To assess for purity, fractions with absorbance at 280 nm were analyzed on a NuPAGE 4-12% Bis-Tris gel, stained with SimplyBlue SafeStain (Invitrogen, LC6060) and imaged on BioRad Gel Doc XR. Fractions with protein of correct weight were concentrated using Millipore Sigma spin-columns (10 kDa cutoff) and buffer-exchanged into TBS. Protein concentrations were determined using a NanoDrop One (Thermo Fisher, ND-ONE-W) using absorbance.

### Standard Calcium Buffers for In Vitro Calibration

Calcium ions are omnipresent, which complicates the knowing the exact amount of ’free calcium’ in solutions. To address this issue, we used a pH sensitive chelator buffer to titrate accurate calcium levels to achieve chelator saturation. We created two buffers, one with calcium titrated in (‘Calcium-Chelator’) and one without any additional calcium (‘Chelator Only’). Dilutions of Calcium-Chelator and Chelator-Only buffers made possible by incrementally increasing levels of ‘free calcium’ in the solution. The buffer was 100 mM nitrilotriacetic acid (NTA), 50 mM MOPS, 100 mM KCl with the pH at 7.2 (KOH to adjust). All pH measurements were done on Orion A2II, calibrated with the supplied buffers. Free Calcium levels were calculated with the calcium-electrode PerfectION (Mettler Toledo, 51344703) and SevenCompact Meter (Mettler Toledo, 300192028).

### Generating and Calculating Curve Fit for In Vitro Calibration

Fluorescence measurements were conducted by adding 1.2 µL of protein to 100 µL of each calcium buffer, achieving a final protein concentration of approximately 1.2 µM. These samples were analyzed by measuring fluorescence on a Tecan Plate Reader Infinite M1000 Pro, with settings adjusted to an excitation wavelength of 485 nm, emission wavelength of 510 nm, a bandwidth of 5 nm, and a gain setting of 80 at a controlled temperature of 37 °C. The resulting fluorescence data were analyzed with GraphPad Prism software, employing the ’Specific Binding with Hill Slope’ model for curve fitting.

### In Cell Calibration of Probes

Ca^2+^ binding curves were determined using cells permeabilized with 25 µM digitonin for 3 minutes at 37 °C. Cells were perfused with solutions that were mixes of 10 mM CaNTA, 100 mM KCl and 50 mM MOPS and 10 mM NTA, 100 mM KCl, 50 mM MOPS. Probes were saturated with a 10 mM CaCl_2_, 100 mM KCl, 50 mM MOPS solution. All calibrations were performed at 37 °C and pH 7.2. Ca^2+^ concentrations for each solution were calculated using (Dweck et al., 2005). To determine the binding properties, fluorescence values were fitted to the generalized Hill equation.

### Immunohistochemistry of hippocampal neurons

Hippocampal cultures transfected with the hDLXI56i-eGFP (or hDLXI56i-mSynaptophysin-pHluorin in some cases) were fixed for 10 minutes in 4% PFA. Following three, five-minute washes with PBS, cultures were placed in blocking buffer (1% BSA, 10% goat serum. 0.3% Triton X-100 in PBS) for 90 minutes. This blocking solution was diluted 1:10 in PBS to use as a staining buffer. Primary antibody was added to the staining buffer and left on the cells overnight at 4 °C. Samples were washed three times for 5 minutes each time in PBS. Secondary antibody was added in staining buffer for 1 hr. Samples were again washed three times for five minutes each in PBS, following mounting in ProLong Diamond. Images were acquired at the same imaging conditions. Primary antibodies and dilutions were 1:500 anti-GFP chicken polyclonal antibody (Invitrogen A10262) and 1:500 anti-GAD2 mouse monoclonal purified IgG antibody (Synaptic Systems 198111). Secondary antibodies and dilutions were 1:500 goat anti-chicken IgY (H+L) secondary antibody, AlexaFluor 488 conjugate (Invitrogen A11039) and 1:500 goat anti-mouse IgY (H+L) secondary antibody, AlexaFluor 568 conjugate (Invitrogen A11029).

### Image Analysis

Images were analyzed using ImageJ and Time Series Analyzer V3 plugin. For two color experiments, images were aligned using the TurboReg plugin (Thevenaz et al., 1998). For ER-Halo-GCaMP-150 experiments in somas, Regions of Interest (ROIs) were drawn around the soma. For axonal regions, the axon was traced using the Halo-JF dye channel. A local background correction was performed using regions next to the axon as the background. All fittings and statistical analysis were performed in GraphPad Prism 8 or 9.

For identification of GAD positive cells from immunohistochemistry, a ROI was placed over the soma of cells that showed GFP staining. If fluorescence was greater than 2X the average fluorescence value for a GAD negative cell in the same dish, the cell was considered GAD positive.

### Ratiometric conversion to Ca^2+^

To report absolute values of Ca^2+^ the ratio was converted to [Ca^2+^] using in cell Ca^2+^ binding curves (*Calibration of Probes*). Fitting was performed using a modified Hill Equation:

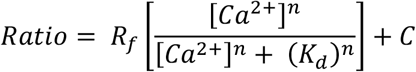

where R_f_ is the dynamic range (F_max_/F_min_), [Ca^2+^] is the concentration of Ca^2+^, K_d_ is the dissociation constant, n is the Hill coefficient, ratio is F_GCaMP_/F_JF dye_, and C is a constant accounting for the fluorescence of the indicator in the presence of 0 Ca^2+^. The above equation can be rearranged to calculate the [Ca^2+^]_ER_ from the ratio:

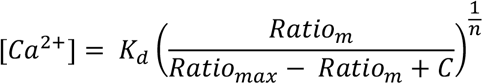

where Ratio_m_ is the measured Ratio (F_GCaMP_/F_JF dye_), Ratio_max_ is the maximum ratio as measured from the calibration curves and all other parameters are as previously described. Conversion from ratio to [Ca^2+^] was calculated using the above equation.

### Statistics

Error bars are reported as ± SEM. Box and whisker plots are Median (Line), 25-75 percentile (box), range (whiskers). Statistical analysis was performed as marked in figure legends. For all analysis p values < 0.05 were considered significant and marked as one asterisk, whereas p < 0.01, p < 0.001 and p < 0.0001 were marked with two, three or four asterisks respectively. Samples were checked for normality and If not normal a non-parametric test was used. The sample number (n) refers to the number of cells. Investigator was not blind to conditions. No estimate of the necessary sample size was performed before experiments.

## Acknowledgements

This work was supported by a grant from the NIH (2R01NS036942) to T.A.R. and a Medical Scientist Training Program grant from the National Institute of General Medical Sciences of the NIH under award number: T32GM007739 to the Weill Cornell/Rockefeller/Sloan Kettering Tri-Institutional MD-PhD Program. We thank Julia Marrs, Lucia A. Saidenberg, Johnny Ye and Hayoung Lee for technical assistance. We thank the members of the Ryan lab for helpful discussion, especially Jaime de Juan-Sanz for generating the ER mRuby3-GCaMP6-210 construct. We thank Luke Lavis for providing the JF_635_ dye. The pKanCMV-mRuby3-18aa-actin was a gift from Michael Lin and the CN1851-rAAV-hI56i-minBglobin-iCre-4×2C-WPRE3-BGHpA was a gift from The Allen Institute for Brain Science & BRAIN Armamentarium & Jonathan Ting.

## Competing Interests

The authors declare no competing interests.

## Author Contributions

Experiments were designed by R.J.F. and T.A.R. T.H. performed the *in vitro* characterization of the ER-GCaMPs under the supervision of T.B. with input from R.J.F and T.A.R. K.G.B. performed the validation of the hDLXI56ipromoter and cloned all hDLXI56i containing constructs. M.B.H performed the CaMKII validation experiments. R.J.F performed all other experiments. The manuscript was written by R.J.F and T.A.R. with input from all the authors.

## Notes

### Competing Interest Statement

The authors have declared no competing interest.

### Summary of Updates

typo in author name

